# LinkExplorer: Predicting, explaining and exploring links in large biomedical knowledge graphs

**DOI:** 10.1101/2022.01.09.475537

**Authors:** Simon Ott, Adriano Barbosa-Silva, Matthias Samwald

**Affiliations:** Institute of Artificial Intelligence, Medical University of Vienna, Vienna, 1090, Austria

## Abstract

**Summary:** Machine learning algorithms for link prediction can be valuable tools for hypothesis generation. However, many current algorithms are black boxes or lack good user interfaces that could facilitate insight into why predictions are made. We present LinkExplorer, a software suite for predicting, explaining and exploring links in large biomedical knowledge graphs. LinkExplorer integrates our novel, rule-based link prediction engine SAFRAN, which was recently shown to outcompete other explainable algorithms and established black box algorithms. Here, we demonstrate highly competitive evaluation results of our algorithm on multiple large biomedical knowledge graphs, and release a web interface that allows for interactive and intuitive exploration of predicted links and their explanations.

**Availability and Implementation:** A publicly hosted instance, source code and further documentation can be found at https://github.com/OpenBioLink/Explorer.

**Supplementary information:** Supplementary data are available at *Bioinformatics* online.

## 1 Introduction

Link prediction is a field in graph-based machine learning that aims to predict novel relationships between entities. When applied to biomedical knowledge graphs, link prediction can be a versatile and powerful method of hypothesis generation, e.g. for drug discovery (Abbas *et al*., 2021). A vast array of machine learning link prediction algorithms emerged over the last decade. Most of these are black boxes such as embedding- or deep learning based algorithms, where the rationale for a prediction is difficult or impossible to explain. Explainability, however, is important for facilitating scientific insight, judging the plausibility of predictions and improving acceptance by end-users (Adadi and Berrada, 2018).

Previous work on explainable link prediction (Meilicke *et al*., 2019) has limitations that decrease its utility for real-world applications involving large biomedical knowledge graphs, such as lacking scalability, limited predictive performance, or lack of end-user interfaces for viewing and understanding predictions and their explanations.

To address these issues, we release the LinkExplorer software suite for predicting, exploring and explaining links in large knowledge graphs. It integrates SAFRAN (Ott *et al*., 2021), a novel, state-of-the-art rule-based link prediction algorithm that our group developed. SAFRAN is highly scalable, outperforms other explainable link prediction methods on standard link prediction benchmarks, narrows the performance gap between explainable and black-box algorithms, and produces highly condensed and transparent explanations (Ott *et al*., 2021). The LinkExplorer suite offers a web-based interface for navigating existing links between entities and relations together with predicted links and their explanations generated by SAFRAN, thereby enabling unprecedented insight into the predictions of a state-of-the-art link prediction algorithm. We also report highly competitive evaluation results of our explainable link prediction algorithm on several large-scale biomedical knowledge graphs.

## 2 Software architecture and performance

In the following we provide a brief overview of the SAFRAN and AnyBURL algorithms. For a detailed description of the algorithms we refer to (Ott *et al*., 2021).

AnyBURL (Meilicke *et al*., 2019) is a walk-based (bottom-up) method based on the idea that sampled paths (random walks) in a knowledge graph are examples of very specific rules and thus can be transformed into more general logical first-order Horn clauses. In each iteration of the rule mining algorithm, AnyBURL samples paths and generalizes them into rules of three predefined types. Examples of rules generalized from a sampled path can be seen in Table 1. Furthermore AnyBURL calculates a confidence for each rule, the probability that an entity predicted by the rule is correct. This confidence is the relative proportion of correctly predicted entities in all predicted entities by a rule when applied to the training set. In order to correct for unseen entities a constant *p*_*c*_ *>* 0 is added to the denominator. As can be seen in Figure 1, LinkExplorer displays the confidence, number of correctly predicted and number of all predicted entities of a rule. In the example of Figure 1 *p*_*c*_ was set to 5.

**Table 1.**
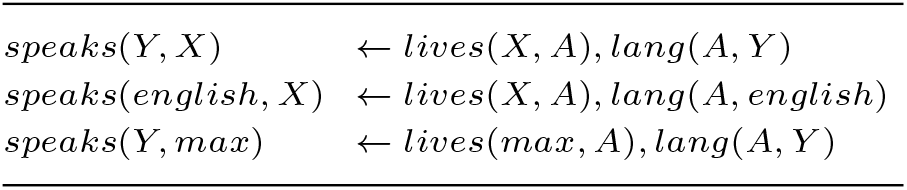
Examples of rules that can be generalized from the sampled path *speaks*(*max, english*) ← *lives*(*max, uk*), *lang*(*uk, english*)

|  |  |
| --- | --- |
| $speaks(Y, X)$ | $\leftarrow lives(X, A), lang(A, Y)$ |
| $speaks(english, X)$ | $\leftarrow lives(X, A), lang(A, english)$ |
| $speaks(Y, max)$ | $\leftarrow lives(max, A), lang(A, Y)$ |

**Fig. 1.**
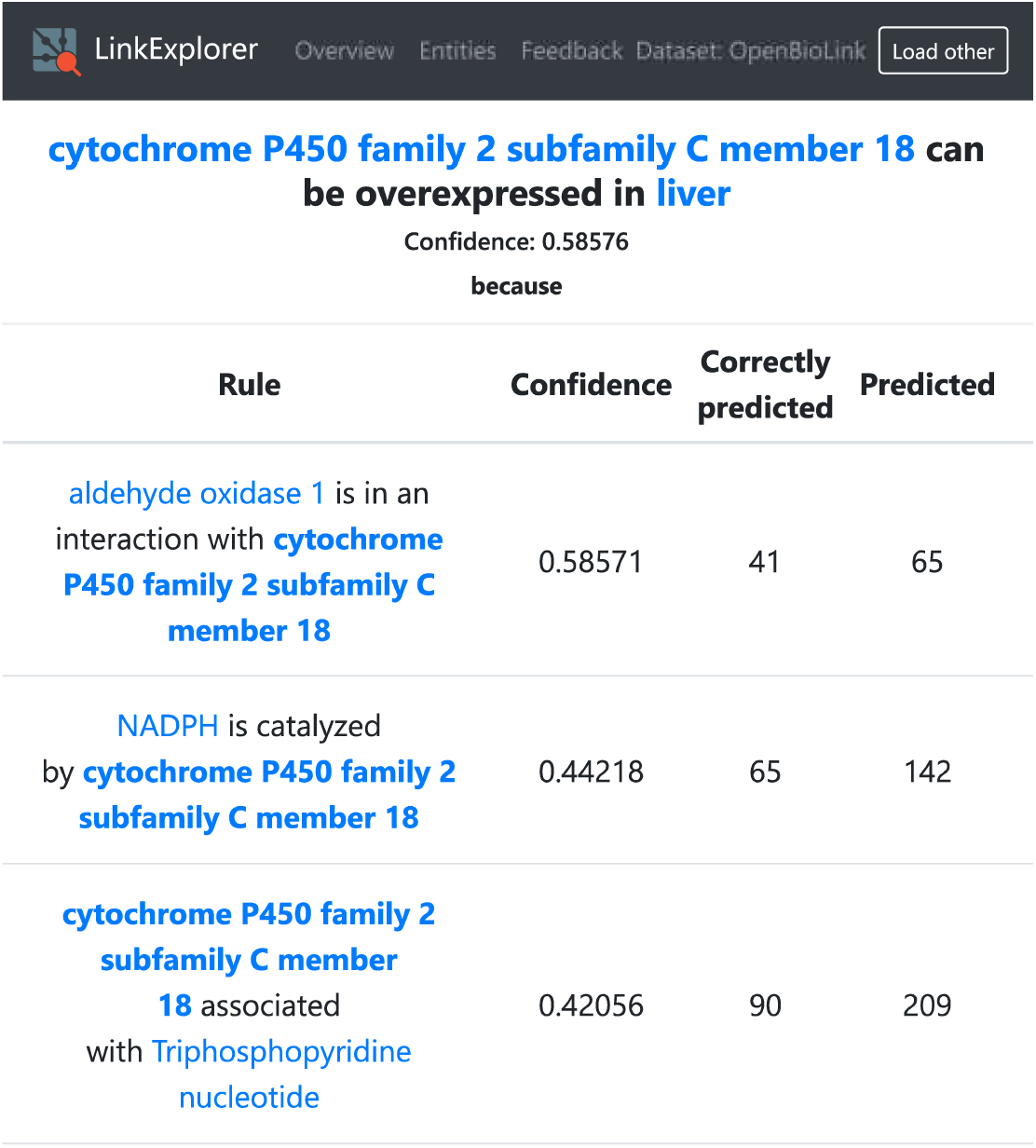
Structured view of rules that triggered a prediction.

The rule-application framework SAFRAN takes rules generated by AnyBURL as input, and improves upon AnyBURL’s rule application method in two important ways. First, SAFRAN was engineered to scale to large knowledge bases, which is essential for many biomedical applications. Second, SAFRAN introduces an algorithm that identifies and clusters redundant rules, i.e. rules that are structurally different but have highly correlated predictions. The predictions of rule clusters can then be aggregated with a noisy-or operation, which would not work well when redundancies are not accounted for. The noisy-or aggregation makes it possible to combine predictions of different rules / rule clusters in a meaningful way and improves predictive accuracy. Furthermore, rule clustering makes generated explanations easier to understand. Each prediction made by SAFRAN can be explained in terms of the rules / rule clusters and instantiations that triggered it. A screenshot of LinkExplorer providing explanations for the prediction *cytochrome P450 family 2 subfamily C member 18 can be overexpressed in liver* is shown in Figure 1. LinkExplorer uses RDF* (Hartig and Thompson, 2019) graphs to store nodes and edges of knowledge graphs which are served via a SPARQL* endpoint. Furthermore these graphs include metadata of entities such as labels, descriptions and types. In addition to loading knowledge graphs hosted on the server, users can load custom knowledge graphs by specifying external SPARQL* endpoints.

The user interface of LinkExplorer was realized using a client-server architecture and is implemented in Javascript and HTML5. The client was created using the ReactJS framework and the server-sided Javascript components are served using NodeJS. The server is used to provide predictions and explanations which are retrieved from SQLite databases generated by SAFRAN. LinkExplorer provides explanation files for all hosted knowledge graphs and allows users to load local prediction files.

We make a public LinkExplorer instance available that hosts three large biomedical knowledge graphs: OpenBioLink (Breit *et al*., 2020), Hetionet (Himmelstein *et al*., 2017) and PheKnowLator (Callahan *et al*., 2020). Additionally, the instance contains two widely-used general-domain evaluation datasets: WN18RR (Dettmers *et al*., 2018) and YAGO3-10 (Dettmers *et al*., 2018). As there is no publicly available dataset split of Hetionet and PheKnowLator, they were randomly split using a 90-5-5 split ratio per relation. Prediction files were generated by SAFRAN using rules that were learned by AnyBURL for 1000 seconds using 22 threads of a machine with 24 physical (48 logical)Intel(R) Xeon(R) CPU E5-2650 v4 @ 2.20GHz cores and 264 GB of RAM. Table 2 shows a comparison of the performance of LinkExplorer/SAFRAN with AnyBURL and established black-box models RESCAL (Nickel *et al*., 2011), TransE (Bordes *et al*., 2013), DistMult (Yang *et al*., 2015), ComplEx (Trouillon *et al*., 2016), ConvE (Dettmers *et al*., 2018), RotatE (Sun *et al*., 2019) for the task of link prediction on the before mentioned biomedical datasets and ogbl-biokg (Hu *et al*., 2020). For training and extensive hyperparameter optimization of the latent models we used the PyTorch-based LibKGE framework (Broscheit *et al*., 2020) and use a similar hyperparameter search space as in (Ruffinelli *et al*., 2020). Data on the conducted hyperparameter search can be found in the supplemental material.

**Table 2.**
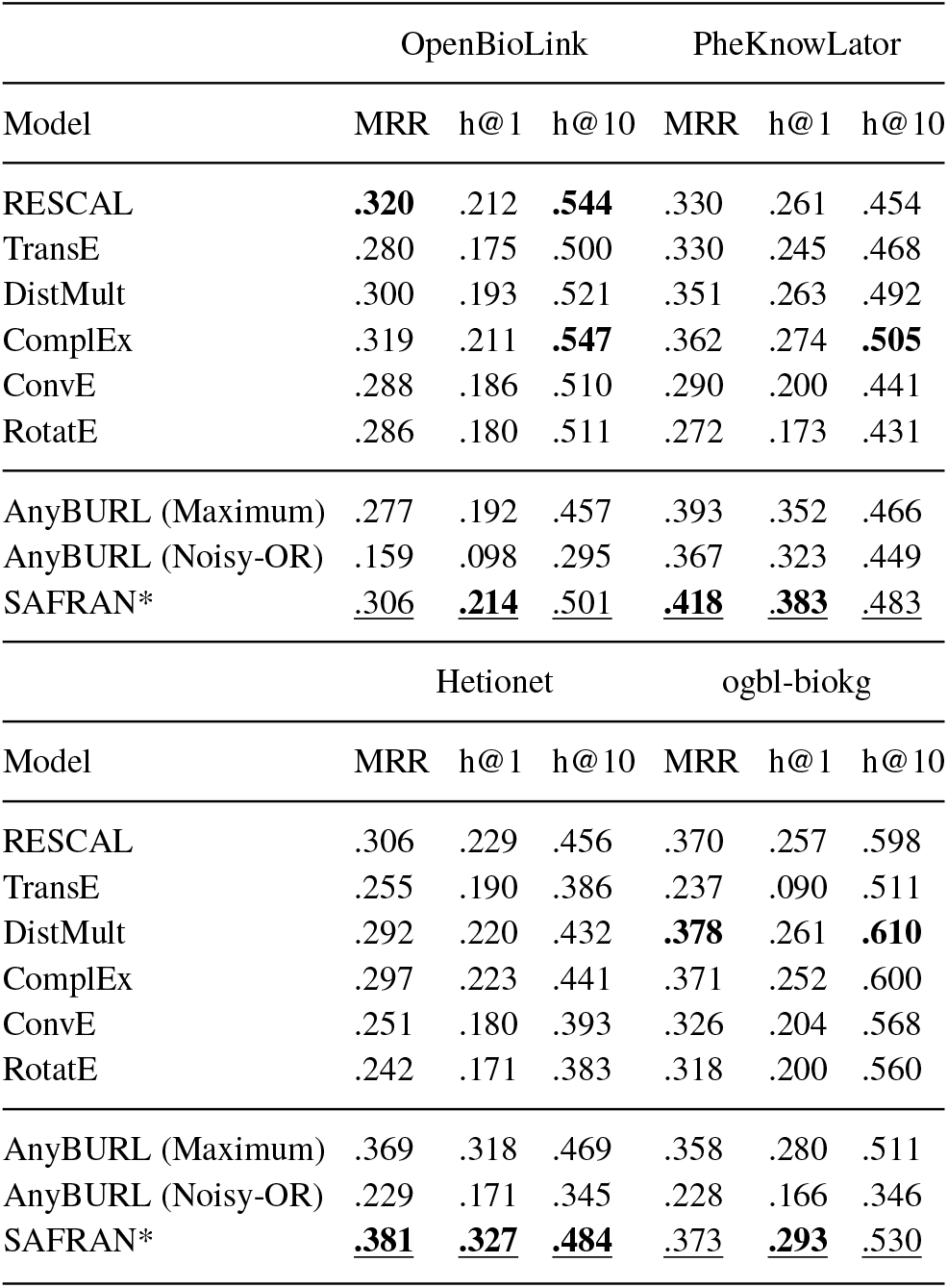
Comparison of the fully explainable algorithms SAFRAN and AnyBURL with several established latent models on the datasets OpenBioLink, PheKnowLator, Hetionet and ogbl-biokg. SAFRAN (denoted with *) is our approach. The best overall results are marked in bold and the best explainable results are underlined.

The evaluation protocol used to evaluate all models link prediction task is as follows: Given a test triple (*h, r, t*) we construct two queries (*h, r*, ?) and (?, *r, t*). For each query, a model scores the likelihood of every entity present in the dataset to act as a substitution for ?. According to the ranked list of scored entities, we report the filtered mean reciprocal rank (MRR), filtered hits@1 and filtered hits@10. MRR is the average reciprocal rank of the correct entity over all queries. Hits@k is the relative proportion of predictions having a rank greater or equal than k. We follow the procedure for filtering known triples first described in (Bordes *et al*., 2013), where triples present in the training, testing and validation sets are removed from the set of predicted triples, with the exception of the correct triple. As symbolic approaches do not scale very well, we calculate the MRR for AnyBURL and SAFRAN based on the top-1000 predicted entities for a query, which results in a result that is marginally less or equal to the actual MRR. All results reported were evaluated according to the average policy Rossi *et al*. (2021) for dealing with same score predictions, where a list of entities is first ranked via their individual score, entities within groups of same score entities are however assigned the average rank of their respective group.

LinkExplorer/SAFRAN consistently improves upon the already highly competitive explainable link prediction algorithm AnyBURL, and closely matches or outperforms black-box models.

## 3 Discussion and future work

LinkExplorer demonstrates that high predictive performance can be achieved while retaining a high level of model explainability and transparency, attributes that are very desirable for biomedical hypothesis generation. Future work should focus on conducting empirical user studies of explanation methods to quantify and understand their utility in real-world biomedical research settings.

## References

Abbas, K., Abbasi, A., Dong, S., Niu, L., Yu, L., Chen, B., Cai, S.-M., and Hasan, Q. (2021). Application of network link prediction in drug discovery. BMC Bioinformatics, 22(1), 187.

Adadi, A. and Berrada, M. (2018). Peeking inside the black-box: A survey on explainable artificial intelligence (xai). IEEE Access, 6, 52138–52160.

Bordes, A., Usunier, N., Garcia-Duran, A., Weston, J., and Yakhnenko, O. (2013). Translating embeddings for modeling multi-relational data.

Breit, A., Ott, S., Agibetov, A., and Samwald, M. (2020). OpenBioLink: a benchmarking framework for large-scale biomedical link prediction. Bioinformatics, 36(13), 4097–4098.

Broscheit, S., Ruffinelli, D., Kochsiek, A., Betz, P., and Gemulla, R. (2020). LibKGE - A knowledge graph embedding library for reproducible research. In Proceedings of the 2020 Conference on Empirical Methods in Natural Language Processing: System Demonstrations, pages 165–174.

Callahan, T. J., Tripodi, I. J., Hunter, L. E., and Baumgartner, W. A. (2020). A framework for automated construction of heterogeneous large-scale biomedical knowledge graphs. bioRxiv.

Dettmers, T., Minervini, P., Stenetorp, P., and Riedel, S. (2018). Convolutional 2d knowledge graph embeddings.In Thirty-Second AAAI Conference on Artificial Intelligence.

Hartig, O. and Thompson, B. (2019). Foundations of an alternative approach to reification in rdf.

Himmelstein, D. S., Lizee, A., Hessler, C., Brueggeman, L., Chen, S. L., Hadley, D., Green, A., Khankhanian, P., and Baranzini, S. E. (2017). Systematic integration of biomedical knowledge prioritizes drugs for repurposing. eLife, 6, e26726.

Hu, W., Fey, M., Zitnik, M., Dong, Y., Ren, H., Liu, B., Catasta, M., and Leskovec, J. (2020). Open graph benchmark: Datasets for machine learning on graphs. In Advances in Neural Information Processing Systems, volume 33, pages 22118–22133. Curran Associates, Inc.

Meilicke, C., Chekol, M. W., Ruffinelli, D., and Stuckenschmidt, H. (2019). Anytime bottom-up rule learning for knowledge graph completion. In Proceedings of the Twenty-Eighth International Joint Conference on Artificial Intelligence, IJCAI-19, pages 3137–3143. International Joint Conferences on Artificial Intelligence Organization.

Nickel, M., Tresp, V., and Kriegel, H.-P. (2011). A three-way model for collective learning on multi-relational data. In Proceedings of the 28th International Conference on International Conference on Machine Learning, ICML’11, page 809–816, Madison, WI, USA. Omnipress.

Ott, S., Meilicke, C., and Samwald, M. (2021). SAFRAN: An interpretable, rule-based link prediction method outperforming embedding models. In 3rd Conference on Automated Knowledge Base Construction.

Rossi, A., Barbosa, D., Firmani, D., Matinata, A., and Merialdo, P. (2021). Knowledge graph embedding for link prediction: A comparative analysis. ACM Trans. Knowl. Discov. Data, 15(2).

Ruffinelli, D., Broscheit, S., and Gemulla, R. (2020). You CAN teach an old dog new tricks! on training knowledge graph embeddings. In International Conference on Learning Representations.

Sun, Z., Deng, Z.-H., Nie, J.-Y., and Tang, J. (2019). Rotate: Knowledge graph embedding by relational rotation in complex space. In International Conference on Learning Representations.

Trouillon, T., Welbl, J., Riedel, S., Gaussier, E., and Bouchard, G. (2016). Complex embeddings for simple link prediction. In M. F. Balcan and K. Q. Weinberger, editors, Proceedings of The 33rd International Conference on Machine Learning, volume 48 of Proceedings of Machine Learning Research, pages 2071–2080, New York, New York, USA. PMLR.

Yang, B., Yih, S. W.-t., He, X., Gao, J., and Deng, L. (2015). Embedding entities and relations for learning and inference in knowledge bases. In Proceedings of the International Conference on Learning Representations (ICLR) 2015.

