## Supplementary material for "LinkExplorer: Predicting, explaining and exploring links in large biomedical knowledge graphs"

Supplementary data

### 1 Statistics on datasets

| | Entities | Relations | Training | Testing | Validation |
|---|---|---|---|---|---|
| OpenBioLink | 184,635 | 28 | 4,192,002 | 183,009 | 180,964 |
| PheKnowLator | 737,556 | 294 | 4,952,471 | 274,497 | 260,775 |
| Hetionet | 45,158 | 24 | 2,030,777 | 112,524 | 106,896 |
| ogbl-biokg | 93,773 | 51 | 4,762,678 | 162,870 | 162,886 |

Table 1: Dataset statistics: Number of entities, relations and triples of each dataset.

| | M-1 | 1-N | M-N | 1-1 |
|---|---|---|---|---|
| OpenBioLink | 2 | - | 25 | 1 |
| PheKnowLator | 63 | 6 | 19 | 206 |
| Hetionet | - | 3 | 21 | - |
| ogbl-biokg | - | - | 51 | - |

Table 2: Number of relation types within each dataset. We use the same approach for classifying relationships as in [Bordes et al., 2013], where four different relationship type classes were used: 1-1, 1-many, many-1, many-many. A relationship is 1-1 if there is at most one tail for each head in the training set and vice versa. A relationship is classified as 1-many if there appear multiple tails with a head, many-1 if there appear multiple heads with a tail and many-many if multiple heads appear with multiple tails. Classes were determined by calculating the average number of tails t (heads h) that appear given a pair (h, r) ((t, r)) in the training set, A threshold of 1.5 average entities was used above which the argument is labeled as many.

| Type | Relation | # Avg. head | # Avg. tail |
|---|---|---|---|
| 1-N | Disease - upregulates - Gene | 1.35 | 158.55 |
| | Disease - downregulates - Gene | 1.29 | 159.95 |
| | Pharmacologic Class - includes - Compound | 1.37 | 2.83 |
| M-N | Gene - interacts - Gene | 9.66 | 14.28 |
| | Anatomy - expresses - Gene | 26.34 | 1979.5 |
| | Disease - presents - Symptom | 7.37 | 22.77 |

Table 3: Examples of relationships of type 1-N and type M-N of Hetionet and their average number of heads per tail and average number of tails per head which lead to their classification.

| Type | Relation | # Avg. head | # Avg. tail |
|---|---|---|---|
| M-1 | IS_A | 4.23 | 1.48 |
| | PART_OF | 2.5 | 1.07 |
| M-N | GENE_UNDEREXPRESSED_ANATOMY | 1202.23 | 7.28 |
| | GENE_PHENOTYPE | 19.54 | 34.39 |
| | DRUG_ACTIVATION_GENE | 4.98 | 3.78 |
| 1-1 | GENE_EXPRESSION_GENE | 1.11 | 1.08 |

Table 4: Examples of relations of different types appearing in OpenBioLink and their average number of heads per tail and average number of tails per head which lead to their classification.

#### 2 Results for FB15K-237

| | | | FB15K-237 | | |
|---|---|---|---|---|---|
| | Approach | | MRR | hits@1 | hits@10 |
| Latent | RESCAL | ⋆ | .357 | .263 | .541 |
| | TransE | ⋆ | .313 | .221 | .497 |
| | DistMult | ⋆ | .343 | .250 | .531 |
| | ComplEx | ⋆ | .348 | .253 | .534 |
| | ConvE | ⋆ | .339 | .248 | .521 |
| | RotatE | ¶ | .336 | .238 | .531 |
| | TuckER | ¶ | .352 | .259 | .536 |
| | HAKE | ▽ | .346 | .250 | **.542** |
| Interpretable | C-NN | △ | .296 | .222 | .446 |
| | DRUM | ‡ | *.343*† | *.255*† | *.516*† |
| | Neural LP | ◊ | *.240*† | | *.362*† |
| | GPFL | ◊ | .322 | .247 | .504 |
| | AMIE+ | ♣ | | .174 | .409 |
| | RuleN | ♣ | | .182 | .420 |
| | RLvLR | ♯ | .240 | | .393 |
| | SAFRAN* | | **.389** | **.298** | .537 |

Table 5: MRR, Hits@1, Hits@10 results for FB15K-237. Best results for each metric and dataset are marked in bold. *Denotes our approach. †Results were evaluated with top policy for dealing with same score entities and are not directly comparable to other approaches. Results marked with ⋆ are from [Ruffinelli et al., 2020], ¶ from [Rossi et al., 2021], △ from [Ferré, 2020], ‡ from [Sadeghian et al., 2019], ◊ from [Gu et al., 2020], ♣ from [Meilicke et al., 2019], ♯ from [Omran et al., 2018] and ▽ from [Zhang et al., 2020].

### 3 Hyperparameter search of embedding models

| Hyperparameter | Range |
|---|---|
| Embedding size | 128 |
| Training type | NegSamp |
| # head samples | $[0, 10000]$ |
| # tail samples | $[0, 10000]$ |
| Loss | CE |
| $L_p$ norm (TransE) | $\{1, 2\}$ |
| Optimizer | Adagrad |
| Batch size | 1024 |
| Learning rate | $[3 * 10^{-4}, 1]$, log scale |
| LR scheduler patience | $[0, 10]$ |
| $L_p$ regularization | $\{1, 2, 3, \text{None}\}$ |
| Entity emb. weight | $[10^{-20}, 10^{-1}]$ |
| Relation emb. weight | $[10^{-20}, 10^{-1}]$ |
| Frequency weighting | True |
| Embedding normalization (TransE) | |
| Entity | $\{\text{True}, \text{False}\}$ |
| Relation | $\{\text{True}, \text{False}\}$ |
| Dropout | |
| Entity embedding | $[0.0, 0.5]$ |
| Relation embedding | $[0.0, 0.5]$ |
| Feature map (ConvE) | $[0.0, 0.5]$ |
| Projection (ConvE) | $[0.0, 0.5]$ |
| Embedding initialization | $\{\text{Normal}, \text{Unif}, \text{XvNorm}, \text{XvUnif}\}$ |
| Std. deviation (normal) | $[10^{-5}, 1.0]$, log scale |
| Interval (Unif) | $[-1.0, 1.0]$ |
| Gain (XvNorm) | 1.0 |
| Gain (XvUnif) | 1.0 |

Table 6: Hyperparameter search space. We follow the naming conventions and ranges given by [Ruffinelli et al., 2020], however fixed some values due to the size of the datasets (training type: negative sampling, embedding size: 128, batch size: 1024, optimizer: Adagrad, regularization: weighted). For the meaning of the parameters, we refer also to this publication. We selected the best configuration based on the MRR of the validation set after running 30 pseudo-random trials for 20 epochs. We retrained the configuration that performed best for a maximum 400 epochs (with an early stopping patience of 10).

| | | RESCAL | TransE | DistMult | ComplEx | ConvE | RotatE |
|---|---|---|---|---|---|---|---|
| *Hetionet* | Embedding size | 128 | 128 | 128 | 128 | 128 | 128 |
| | Training type | NegSamp | NegSamp | NegSamp | NegSamp | NegSamp | NegSamp |
| | Reciprocal | No | No | No | No | Yes | No |
| | No. subject samples (NegSamp) | 7336 | 2539 | 7336 | 7336 | 7986 | 2687 |
| | No. object samples (NegSamp) | 5160 | 7092 | 5160 | 5160 | 1325 | 2804 |
| | Label Smoothing (KvsAll) | – | – | – | – | – | – |
| | Loss | CE | CE | CE | CE | CE | CE |
| | Margin (MR) | – | – | – | – | – | – |
| | $L_p$-norm (TransE) | – | L1 | – | – | – | – |
| | Optimizer | Adagrad | Adagrad | Adagrad | Adagrad | Adagrad | Adagrad |
| | Batch size | 1024 | 1024 | 1024 | 1024 | 1024 | 1024 |
| | Learning rate | 0.11374 | 0.08972 | 0.11374 | 0.11374 | 0.02525 | 0.04883 |
| | Scheduler patience | 8 | 4 | 8 | 8 | 3 | 6 |
| | $L_p$ regularization | L3 | L3 | L3 | L3 | L2 | L1 |
| | Entity emb. weight | $2.61^{-20}$ | $3.32^{-17}$ | $2.61^{-20}$ | $2.61^{-20}$ | $2.26^{-15}$ | $5.55^{-16}$ |
| | Relation emb. weight | $2.00^{-20}$ | $8.04^{-18}$ | $2.00^{-20}$ | $2.00^{-20}$ | $1.61^{-06}$ | $2.28^{-08}$ |
| | Frequency weighting | Yes | Yes | Yes | Yes | Yes | Yes |
| | Embedding normalization (TransE) | | | | | | |
| | Entity | – | L2 | – | – | – | – |
| | Relation | – | No | – | – | – | – |
| | Dropout | | | | | | |
| | Entity embedding | 0.00 | 0.00 | 0.00 | 0.00 | 0.00 | 0.00 |
| | Relation embedding | 0.27 | 0.00 | 0.27 | 0.27 | 0.00 | 0.00 |
| | Projection (ConvE) | – | – | – | – | 0.26 | – |
| | Feature map (ConvE) | – | – | – | – | 0.11 | – |
| | Embedding initialization | XvNorm | Unif. | XvNorm | XvNorm | XvUnif | XvUnif |
| | Std. deviation (Normal) | – | – | – | – | – | – |
| | Interval (Unif) | – | [-0.30, 0.30] | – | – | – | – |
| | Gain (XvNorm) | – | – | – | – | 1.00 | 1.00 |
| | Gain (XvUnif) | 1.00 | – | 1.00 | 1.00 | – | – |

Table 7: Best link prediction hyperparameter configurations for each model on Hetionet

| | | RESCAL | TransE | DistMult | ComplEx | ConvE | RotatE |
|---|---|---|---|---|---|---|---|
| *PheKnowLator* | Embedding size | 128 | 128 | 128 | 128 | 128 | 128 |
| | Training type | NegSamp | NegSamp | NegSamp | NegSamp | NegSamp | NegSamp |
| | Reciprocal | No | No | No | No | Yes | No |
| | No. subject samples (NegSamp) | 8985 | 2539 | 9193 | 9193 | 9281 | 2687 |
| | No. object samples (NegSamp) | 7990 | 7092 | 9096 | 9096 | 9430 | 2804 |
| | Label Smoothing (KvsAll) | – | – | – | – | – | – |
| | Loss | CE | CE | CE | CE | CE | CE |
| | Margin (MR) | – | – | – | – | – | – |
| | $L_p$-norm (TransE) | – | L1 | – | – | – | – |
| | Optimizer | Adagrad | Adagrad | Adagrad | Adagrad | Adagrad | Adagrad |
| | Batch size | 1024 | 1024 | 1024 | 1024 | 1024 | 1024 |
| | Learning rate | 0.10137 | 0.08972 | 0.18781 | 0.18781 | 0.05670 | 0.04883 |
| | Scheduler patience | 1 | 4 | 2 | 2 | 2 | 6 |
| | $L_p$ regularization | L2 | L3 | L2 | L2 | L2 | L1 |
| | Entity emb. weight | $2.33^{-02}$ | $3.32^{-17}$ | $3.76^{-19}$ | $3.76^{-19}$ | $9.03^{-20}$ | $5.55^{-16}$ |
| | Relation emb. weight | $1.90^{-07}$ | $8.04^{-18}$ | $2.00^{-20}$ | $2.00^{-20}$ | $4.34^{-16}$ | $2.28^{-08}$ |
| | Frequency weighting | Yes | Yes | Yes | Yes | Yes | Yes |
| | Embedding normalization (TransE) | | | | | | |
| | Entity | – | L2 | – | – | – | – |
| | Relation | – | No | – | – | – | – |
| | Dropout | | | | | | |
| | Entity embedding | 0.00 | 0.00 | 0.00 | 0.00 | 0.00 | 0.00 |
| | Relation embedding | 0.00 | 0.00 | 0.49 | 0.49 | 0.41 | 0.00 |
| | Projection (ConvE) | – | – | – | – | 0.14 | – |
| | Feature map (ConvE) | – | – | – | – | 0.05 | – |
| | Embedding initialization | XvUnif | Unif. | Normal | Normal | Normal | XvUnif |
| | Std. deviation (Normal) | – | – | 0.00003 | 0.00003 | 0.00003 | – |
| | Interval (Unif) | – | [-0.30, 0.30] | – | – | – | – |
| | Gain (XvNorm) | 1.00 | – | – | – | – | 1.00 |
| | Gain (XvUnif) | – | – | – | – | – | – |

Table 8: Best link prediction hyperparameter configurations for each model on PheKnowLator

| | | RESCAL | TransE | DistMult | ComplEx | ConvE | RotatE |
|---|---|---|---|---|---|---|---|
| *ogbl-biokg* | Embedding size | 128 | 128 | 128 | 128 | 128 | 128 |
| | Training type | NegSamp | NegSamp | NegSamp | NegSamp | NegSamp | NegSamp |
| | Reciprocal | No | No | No | No | Yes | No |
| | No. subject samples (NegSamp) | 8985 | 3791 | 7712 | 7712 | 9281 | 8985 |
| | No. object samples (NegSamp) | 7990 | 2876 | 600 | 600 | 9430 | 7990 |
| | Label Smoothing (KvsAll) | – | – | – | – | – | – |
| | Loss | CE | CE | CE | CE | CE | CE |
| | Margin (MR) | – | – | – | – | – | – |
| | $L_p$-norm (TransE) | – | L1 | – | – | – | – |
| | Optimizer | Adagrad | Adagrad | Adagrad | Adagrad | Adagrad | Adagrad |
| | Batch size | 1024 | 1024 | 1024 | 1024 | 1024 | 1024 |
| | Learning rate | 0.10137 | 0.22584 | 0.11374 | 0.11374 | 0.05670 | 0.10137 |
| | Scheduler patience | 1 | 5 | 8 | 8 | 2 | 1 |
| | $L_p$ regularization | L2 | L1 | L2 | L2 | L2 | L2 |
| | Entity emb. weight | $2.33^{-02}$ | $2.03^{-16}$ | $2.61^{-20}$ | $2.61^{-20}$ | $9.03^{-20}$ | $2.33^{-02}$ |
| | Relation emb. weight | $1.90^{-07}$ | $1.75^{-10}$ | $8.48^{-07}$ | $8.48^{-07}$ | $4.34^{-16}$ | $1.90^{-07}$ |
| | Frequency weighting | Yes | Yes | Yes | Yes | Yes | Yes |
| | Embedding normalization (TransE) | | | | | | |
| | Entity | – | No | – | – | – | – |
| | Relation | – | No | – | – | – | – |
| | Dropout | | | | | | |
| | Entity embedding | 0.00 | 0.00 | 0.00 | 0.00 | 0.00 | 0.00 |
| | Relation embedding | 0.00 | 0.00 | 0.35 | 0.35 | 0.41 | 0.00 |
| | Projection (ConvE) | – | – | – | – | 0.14 | – |
| | Feature map (ConvE) | – | – | – | – | 0.05 | – |
| | Embedding initialization | XvUnif | XvUnif | Normal | Normal | Normal | XvUnif |
| | Std. deviation (Normal) | – | – | 0.00564 | 0.00564 | 0.00003 | – |
| | Interval (Unif) | – | – | – | – | – | – |
| | Gain (XvNorm) | 1.00 | 1.00 | – | – | – | 1.00 |
| | Gain (XvUnif) | – | – | – | – | – | – |

Table 9: Best link prediction hyperparameter configurations for each model on ogbl-biokg

| | | RESCAL | TransE | DistMult | ComplEx | ConvE | RotatE |
|---|---|---|---|---|---|---|---|
| *OpenBioLink* | Embedding size | 128 | 128 | 128 | 128 | 128 | 128 |
| | Training type | NegSamp | NegSamp | NegSamp | NegSamp | NegSamp | NegSamp |
| | Reciprocal | No | No | No | No | Yes | No |
| | No. subject samples (NegSamp) | 8985 | 2539 | 8985 | 9193 | 9281 | 8985 |
| | No. object samples (NegSamp) | 7990 | 7092 | 7990 | 9096 | 9430 | 7990 |
| | Label Smoothing (KvsAll) | – | – | – | – | – | – |
| | Loss | CE | CE | CE | CE | CE | CE |
| | Margin (MR) | – | – | – | – | – | – |
| | $L_p$-norm (TransE) | – | L1 | – | – | – | – |
| | Optimizer | Adagrad | Adagrad | Adagrad | Adagrad | Adagrad | Adagrad |
| | Batch size | 1024 | 1024 | 1024 | 1024 | 1024 | 1024 |
| | Learning rate | 0.10137 | 0.08972 | 0.10137 | 0.18781 | 0.05670 | 0.10137 |
| | Scheduler patience | 1 | 4 | 1 | 2 | 2 | 1 |
| | $L_p$ regularization | L2 | L3 | L2 | L2 | L2 | L2 |
| | Entity emb. weight | $2.33^{-02}$ | $3.32^{-17}$ | $2.33^{-02}$ | $3.76^{-19}$ | $9.03^{-20}$ | $2.33^{-02}$ |
| | Relation emb. weight | $1.90^{-07}$ | $8.04^{-18}$ | $1.90^{-07}$ | $2.00^{-20}$ | $4.34^{-16}$ | $1.90^{-07}$ |
| | Frequency weighting | Yes | Yes | Yes | Yes | Yes | Yes |
| | Embedding normalization (TransE) | | | | | | |
| | Entity | – | L2 | – | – | – | – |
| | Relation | – | No | – | – | – | – |
| | Dropout | | | | | | |
| | Entity embedding | 0.00 | 0.00 | 0.00 | 0.00 | 0.00 | 0.00 |
| | Relation embedding | 0.00 | 0.00 | 0.00 | 0.49 | 0.41 | 0.00 |
| | Projection (ConvE) | – | – | – | – | 0.14 | – |
| | Feature map (ConvE) | – | – | – | – | 0.05 | – |
| | Embedding initialization | XvUnif | Unif. | XvUnif | Normal | Normal | XvUnif |
| | Std. deviation (Normal) | – | – | – | 0.00003 | 0.00003 | – |
| | Interval (Unif) | – | [-0.30, 0.30] | – | – | – | – |
| | Gain (XvNorm) | 1.00 | – | 1.00 | – | – | 1.00 |
| | Gain (XvUnif) | – | – | – | – | – | – |

Table 10: Best link prediction hyperparameter configurations for each model on OpenBioLink

### References


Antoine Bordes, Nicolas Usunier, Alberto Garcia-Duran, Jason Weston, and Oksana Yakhnenko. Translating embeddings for modeling multi-relational data. 2013. URL `https://papers.nips.cc/paper/5071-translating-embeddings-for-modeling-multi-relational-dat`

Sébastien Ferré. Application of concepts of neighbours to and knowledge graph completion. *Data Science*, Preprint:1–28, 2020. ISSN 2451-8492. doi: 10.3233/DS-200030. URL `https://doi.org/10.3233/DS-200030`. Preprint.

Yulong Gu, Yu Guan, and Paolo Missier. Towards learning instantiated logical rules from knowledge graphs, 2020.

Christian Meilicke, Melisachew Wudage Chekol, Daniel Ruffinelli, and Heiner Stuckenschmidt. Anytime bottom-up rule learning for knowledge graph completion. In Sarit Kraus, editor, *Proceedings of International Joint Conferences on Artificial Intelligence*, pages 3137–3143, California, aug 2019. ISBN 978-0-9992411-4-1. doi: 10.24963/ijcai.2019/435.

Pouya Ghiasnezhad Omran, Kewen Wang, and Zhe Wang. Scalable rule learning via learning representation. In *Proceedings of the Twenty-Seventh International Joint Conference on Artificial Intelligence, IJCAI-18*, pages 2149–2155. International Joint Conferences on Artificial Intelligence Organization, 7 2018. doi: 10.24963/ijcai.2018/297. URL `https://doi.org/10.24963/ijcai.2018/297`.

Andrea Rossi, Denilson Barbosa, Donatella Firmani, Antonio Matinata, and Paolo Merialdo. Knowledge graph embedding for link prediction: A comparative analysis. *ACM Trans. Knowl. Discov. Data*, 15 (2), January 2021. ISSN 1556-4681. doi: 10.1145/3424672. URL `https://doi.org/10.1145/3424672`.

Daniel Ruffinelli, Samuel Broscheit, and Rainer Gemulla. You can teach an old dog new tricks! on training knowledge graph embeddings. In *International Conference on Learning Representations*, 2020. URL `https://openreview.net/forum?id=BkxSmlBFvr`.

Ali Sadeghian, Mohammadreza Armandpour, Patrick Ding, and Daisy Zhe Wang. Drum: End-to-end differentiable rule mining on knowledge graphs. In H. Wallach, H. Larochelle, A. Beygelzimer, F. d'Alché-Buc, E. Fox, and R. Garnett, editors, *Advances in Neural Information Processing Systems*, volume 32. Curran Associates, Inc., 2019.

Zhanqiu Zhang, Jianyu Cai, Yongdong Zhang, and Jie Wang. Learning hierarchy-aware knowledge graph embeddings for link prediction. *Proceedings of the AAAI Conference on Artificial Intelligence*, 34 (03):3065–3072, Apr. 2020. doi: 10.1609/aaai.v34i03.5701. URL `https://ojs.aaai.org/index.php/AAAI/article/view/5701`.
